# Energy Optimization is a Major Objective in the Real-Time Control of Step Width in Human Walking

**DOI:** 10.1101/506808

**Authors:** Sabrina J. Abram, Jessica C. Selinger, J. Maxwell Donelan

## Abstract

People prefer to move in energetically optimal ways during walking. We have recently found that this preference arises not just through evolution and development, but that the nervous system will continuously optimize step frequency in response to new energetic cost landscapes. Here we test whether energy optimization is also a major objective in the nervous system ‘ s real-time control of step width. To accomplish this, we use a device that can reshape the relationship between step width and energetic cost, shifting the energy optimal width wider than that initially preferred. We find that the nervous system doesn ‘ t spontaneously initiate energy optimization, but instead requires experience with a lower energetic cost step width. After initiating optimization, people converge towards their new energy optimal width within hundreds of steps and update this as their new preferred width, rapidly returning to it when perturbed away. However, energy optimization was incomplete as this new preferred width was slightly narrower than the energetically optimal width. This suggests that the nervous system may determine its preferred width by optimizing energy simultaneously with other objectives such as stability or maneuverability. Collectively, these findings indicate that the nervous systems of able-bodied people continuously optimize energetic cost to determine preferred step width.

## INTRODUCTION

When we walk, we tend to prefer a particular step width and execute this preference with remarkably small variability. For young healthy walkers, preferred step width is approximately 12 cm and varies between steps by less than two centimeters (Donelan et al., 2001; 2004). However, this preference can be influenced by walking context. For example, increases in speed result in decreases in step width and increases in its variability (Stimpson et al., 2018). Another context is stability—preferred width and its variability decrease with external lateral stabilization (Dean et al., 2007; Donelan et al., 2004) and increase with visual field perturbations (Bauby and Kuo, 2000; McAndrew et al., 2010; O’Connor and Kuo, 2009). Step width has also been found to increase with both age and obesity (Brach et al., 2008; Browning and Kram, 2007; Gabell and Nayak, 1984; Owings and Grabiner, 2004a; 2004b; Spyropoulos et al., 1991). One way to understand how the nervous system controls step width is to identify the relative importance of different walking objectives in response to changes in walking contexts.

One well established objective of the nervous system is to optimize metabolic energetic cost. Decades of research show that people naturally prefer to move in energetically optimal ways (Alexander, 1996; Ralston, 1958; Zarrugh et al., 1974). In the case of step width, energy costs are largely determined by step-to-step transition costs, lateral limb swing costs, and active stabilization costs (Donelan et al., 2001; 2002; 2004; Shipman et al., 2002; Shorter et al., 2017). People prefer to walk near the energetically optimal step width, avoiding high step-to-step transition costs at wide widths, as well as high lateral limb swing and active lateral stabilization costs at narrow widths (Donelan et al., 2002; 2001; 2004; Shipman et al., 2002; Shorter et al., 2017). However, although the preferred width is near the energy optimal width in familiar walking conditions, this does not necessarily mean that energy optimality drives the nervous system ‘ s real-time control of width. Instead, the preference may arise from long time-scale processes such as evolution and development (Alexander, 2001; 1996; Rodman and McHenry, 1980; Sockol et al., 2007).

In determining preferred step width, the nervous system likely considers objectives other than energy, such as stability or maneuverability. After all, energy is not preeminent in walking—we must first remain stable and not fall over (Dean et al., 2007; McAndrew Young and Dingwell, 2012), and then we must maneuver through our environment to reach our walking goal (Acasio et al., 2017; Jindrich and Qiao, 2009; Patla et al., 1991; Wu et al., 2015). As one example of an objective function, the nervous system may seek to simultaneously optimize energy, stability, and maneuverability (Huang and Ahmed, 2011). The execution of such a multi-term optimization can vary in a number of ways. First, the terms may not contribute equally to the objective function. That is, they may be independently weighted so that optimizing energy, for example, is treated as a higher priority. Second, the weightings need not be fixed, but may instead depend on walking conditions. For example, the priority for stability may be increased when the consequences of falling are more severe, such as for the elderly (Lord et al., 1999; Rogers et al., 2001; Rogers and Mille, 2003). Third, some objectives may be treated as constraints. For example, a stability constraint may dictate that we walk with widths wider than a particular width. As long as we fulfill this constraint, the nervous system may no longer be concerned about optimizing stability, and instead seek to optimize the other contributors to its objective function. It remains unknown if and how the nervous system represents the real-time control of step width as a multi-term optimization.

The purpose of this study was to test if energy optimization is a major objective in the real-time control of step width. Our lab previously found that people adapt their step frequency to converge on new energy optima, demonstrating that energy is indeed a major objective in the real-time control of some aspects of walking (Selinger et al., 2015). Here we hypothesize that step width is similarly controlled to optimize energy. To test this hypothesis, we shifted the energy optimal width and observed the nervous system ‘ s response. To accomplish this, we built a custom device that applies energetic penalties, in real-time, as a function of measured step width. If optimizing energy is a major objective in the nervous system ‘ s control of step width, we expect that it would adapt its preferred width towards the new energy optimal width.

## METHODS

### Experimental Design

After Simha and Donelan, we built a simple mechatronic system to shift people s energy optimal step width (Simha and Donelan, 2018). In this system, all subjects walked on an instrumented split-belt treadmill at 1.25 m/s (FIT, Bertec Corporation, Columbus, OH, USA). To shift people ‘ s energy optimal width, we commanded the treadmill incline based on the desired energetic penalty for the step width measured from the previous step (Fig. 1a). We implemented this closed loop control of incline based on measured step width using Simulink Real-Time Workshop running at 200 Hz (Simulink Real-Time Workshop, MathWorks Inc., Natick, MA, USA). This controller estimated step width by measuring forces and moments, calculating the left and right centers of pressure, and taking the difference between the lateral centers of pressure for consecutive steps. It first filtered forces and moments using a second-order, low-pass, one-way, digital Butterworth filter (15 Hz cut-off), and then calculated the left and right lateral centers of pressure by dividing the left and right lateral moments by their vertical forces (Verkerke et al., 2005). We identified foot contact events from the characteristic rapid fore-aft translation in ground reaction force center of pressure.

**Fig. 1.**
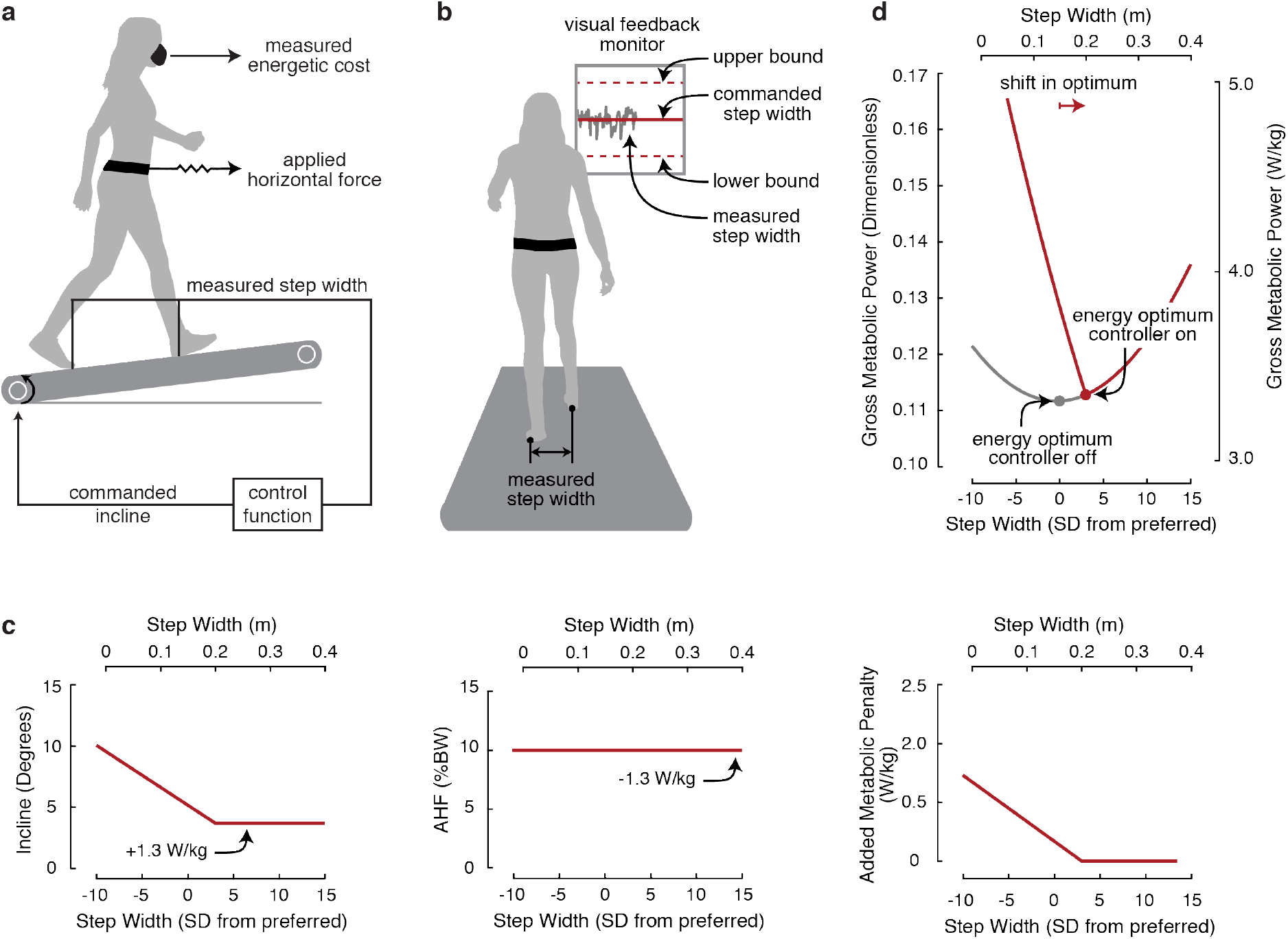
Experimental setup and design. (a) Our real-time controller uses forces and moments measured by the treadmill to identify foot contact events, calculate step width, and then command the appropriate treadmill incline based on the desired energetic penalty for the measured step width. (b) Our real-time visual feedback system allows us to enforce specific widths by instructing subjects to keep their measured width signal within small bounds about the commanded width. (c) We used simulations to design the control function, where the added energetic penalty is high at narrow widths and low at wide widths, by combining literature values for energetic costs of walking at different widths, at different inclines, and with a 10% body weight (BW) applied horizontal force (AHF). A negative step width indicates that the subjects are walking with their steps laterally crossing over one another. (d) We added the energetic cost of the control function to the original cost landscape (grey) to predict the new cost landscape (red) with the energy optimum shifted 3 SD wider than that initially preferred.

We combined the controllable energetic penalty, achieved by manipulating treadmill incline, with a near constant energetic reward, achieved by a forward horizontal force applied to the user. We applied this forward force to disassociate the new energy optimal width from level walking, in the event that the nervous system simply preferred to walk at level inclines, rather than minimize energy. To apply the forward horizontal force, we connected a tensioned cable in series with long rubber tubing to a hip belt worn by the user—the long and compliant rubber tubing allowed for small changes in the walking position on the treadmill without large changes in the applied horizontal force. We adjusted the tension to apply a 10% body weight force, on average, and monitored the applied horizontal force with a load cell mounted on a hip-belt (LCM201, Omega Engineering, Norwalk, CT, USA). When combined with the energetic reward of the forward force, the range of walking incline was from 4.0 ± 1.3° to 8.4 ± 1.2°.

We used this system to reshape the relationship between step width and energetic cost by providing energetic penalties as a function of step width. We define the relationship between gait and total energetic cost as ‘cost landscape ‘ and the relationship between gait and energetic penalties as ‘control function ‘. Similar to Simha and Donelan (Simha and Donelan, 2018), we designed our control function using simulations that combined literature values for energetic costs of walking at different step widths, at different inclines, and with a forward horizontal force (Donelan et al., 2001; Gottschall and Kram, 2003; Margaria, 1968). We added the energetic cost of the control function (Fig. 1c) to the original cost landscape to produce the new cost landscape (Fig. 1d). We designed the control function to produce a new cost landscape with both a clear energetic gradient about each subject’s initial preferred width and a new energy optimal width three standard deviations (SD) wider than each subject s initial preferred. To achieve a clear energetic gradient, we designed an energetic penalty of 2 W/kg at a step width of 3 SD narrower than initial preferred, which decreased with a constant slope to no energetic penalty (0 W/kg) at the new energy optimal width of 3 SD wider than initial preferred. We normalized width by width variability to allow us to distinguish between a shift in width occurring as a result of energy optimization, and a shift occurring by random chance. We chose to shift the energy optimal width wider than that initially preferred in order to be in the opposite direction of people’s tendency to narrow their step width as they become more comfortable walking on a treadmill (Zeni and Higginson, 2010).

### Experimental Protocol

Eight subjects (body mass: 62.6 ± 9.0 kg; height: 165.3 ± 9.5 cm; mean ± SD) participated in the study. All subjects were healthy and exhibited no clinical gait abnormalities. The Simon Fraser University Research Ethics board approved the protocol and participants gave their written, informed consent before completing the experiment.

First, subjects completed a baseline trial to measure their initial preferred width and width variability (Fig. 2a). They did this while walking on the level, without the control of treadmill incline, and without a forward pulling force. We calculated *initial preferred width* as the average width during the final 3 minutes of the 12-minute baseline trial. We calculated each subject s width variability as the standard deviation during the same averaging window. For each subject, we used their initial preferred width and width variability to design the previously described control function. We found that subjects had an average width of 15.4 ± 3.8 cm (mean ± SD) and width variability of 1.5 ± 0.2 cm.

**Fig. 2.**
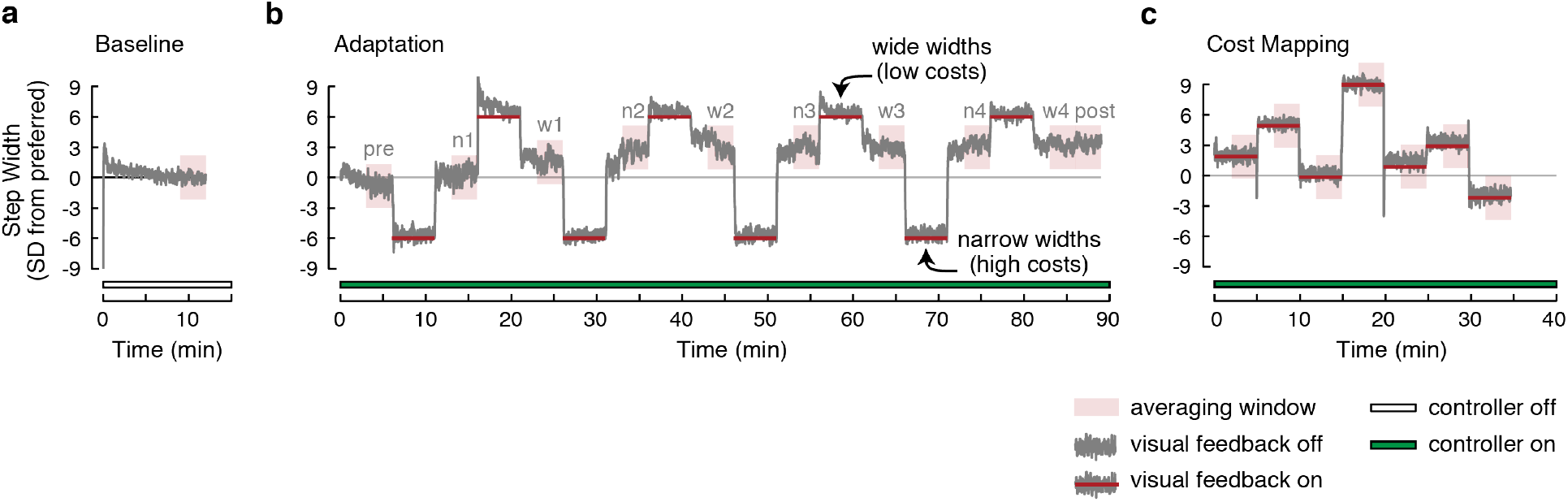
Experimental protocol. Each subject completed all three trials. We averaged measured step width across subjects. (a) First, before we engaged the controller, each subject completed a baseline trial where we determined their preferred width and width variability. (b) Next, each subject completed the experience period where we measured whether they would adapt their preferred width towards the new energy optimal width spontaneously, and when given enforced experience with the new cost landscape through perturbations to different widths. During the periods denoted by the red horizontal lines, we used visual feedback to command either narrow widths (higher costs) or wider widths (lower costs). During the periods without the red horizontal lines, subjects walked at self-selected widths. (c) Lastly, each subject competed a cost mapping trial where we measured their energetic cost at different widths in the new cost landscape. The regions of red shading illustrate the averaging window for steady-state step widths.

Next, we turned the controller on and measured whether subjects adapted their step width towards the new energy optimal width. First, we measured whether subjects would adapt their width spontaneously, calculated as the average width during the final 3 minutes of the first 6-minutes after the controller was turned on (Fig. 2b, pre). Second, we measured whether subjects would adapt their width when given enforced experience with the new cost landscape. This enforced experience consisted of eight perturbations—each perturbation included a 5-minute hold, where subjects were visually-guided to walk with either narrower widths (higher costs) or wider widths (lower costs), followed by a 5-minute release, during which subjects self-selected their widths (Fig. 2b). This visual guidance was given through real-time visual feedback, displayed on a computer monitor, to command target widths (Fig. 1b). The monitor displayed the real-time calculated width, normalized to the commanded width, and instructed subjects to keep the normalized signal close to the commanded width. Subjects successfully matched the commanded width with an average steady-state error of 0.8 ± 0.3 cm. We calculated the change in preferred width at each perturbation as the average width during the final three minutes of the release. We calculated each subject’s *final preferred width* as the average width during the three minutes following the last period of enforced experience (Fig. 2b, post).

Lastly, subjects completed a cost mapping trial where we used visual feedback to enforce steady-state walking at different widths for five minutes each (Fig. 2c). We used respiratory gas analysis (Vmax Encore Metabolic Cart, ViaSys, Conshohocken, PA, USA) to measure rates of oxygen consumption and carbon dioxide production. Subjects walked at widths both about the initial preferred width (−2, 0, +2 SD from initial preferred) and about the final preferred width (−2, 0, +2, +6 SD from final preferred). We calculated metabolic power using the Brockway equation (Brockway, 1987) and determined metabolic power at specific step widths by averaging over the final three minutes of each 5-minute walking period (Selinger and Donelan, 2014).

### Analysis

We compared the new cost landscape to an original cost landscape to verify that we had shifted the energy optimal width wider than that initially preferred. Because we did not collect the original cost landscape in our current experiment, we leveraged existing data collected in different subjects (n=10) (Donelan et al., 2001). For the new cost landscape, we converted step width into units of standard deviations from preferred by first subtracting each subject’s initial preferred width and then dividing by the standard deviation of this width. Because individual step width variability was not measured in the original cost landscape experiments, we used the average standard deviation from our new subjects to convert the original cost landscape into units of standard deviations from preferred (Donelan et al., 2001). For both the original and new cost landscapes, we performed all energetic cost analysis in dimensionless units. We made energetic cost, measured in W/kg, dimensionless using a normalization factor of *g*^3^/^2^*L*^1/2^, where *g* is gravitational acceleration 9.81 ms^−2^ and *L* is leg length calculated as 0.53 m multiplied by the measured height (Drillis et al., 1969). To convert dimensionless units back into W/kg, we divide by the average normalization factor of 29.7 for the original cost landscape and 28.7 for the new cost landscape. We calculated each subject’s energy optimum by first averaging step width and energetic cost over each steady-state walking period, fitting a second-order polynomial, and then calculating the minimum of this polynomial. We calculated the original and new energy optimal widths as the average fitted minima across subjects in the original and new cost landscapes. We calculated the variability in these energy optimal widths as the standard deviation of the fitted minima across subjects. We used a one-tailed paired Student’s t-test to test if the new energy optimal widths were wider than initial preferred widths.

We also tested for adaptation towards the energy optimal width throughout the protocol. We used one-tailed paired Student’s t-tests to determine if each subject’s self-selected width, averaged during the final three minutes of each release, was wider than their initial preferred width. To determine the rate of adaptation, we combined self-selected widths during the release periods after optimization was initiated, averaged across subjects, and then used a first-order exponential to determine the time constant (Fig. 4a). We used two-tailed paired Student s t-tests to test for differences in the final preferred widths compared to the new energy optimal widths, as well as for differences in the costs of these widths. We calculated the cost of each subject’s final preferred width by commanding this width during the cost mapping trial, and the cost of each subject’s energy optimal width as the minimum metabolic power of the fitted new cost landscape. We considered a p-value of 0.05 to be significant in all cases.

## RESULTS

Our control system successfully shifted the energy optimal step width wider than that initially preferred (*p* = 1.3 x 10^−5^; Fig. 3). In the previously collected original cost landscape, the average energy optimal width (0.10 ± 0.05 m; mean±SD) was 1.1 ± 3.5 SD slightly narrower than the original preferred width (0.12 ± 0.01 m) (Donelan et al., 2001). The *original preferred width* is subjects ‘ preferred width from original cost landscape experiments. In our new cost landscape, the average energy optimal width (0.24 ± 0.03 m) was 4.9 ± 1.1 SD wider than the initial preferred width (0.15 ± 0.04 m).

**Fig. 3.**
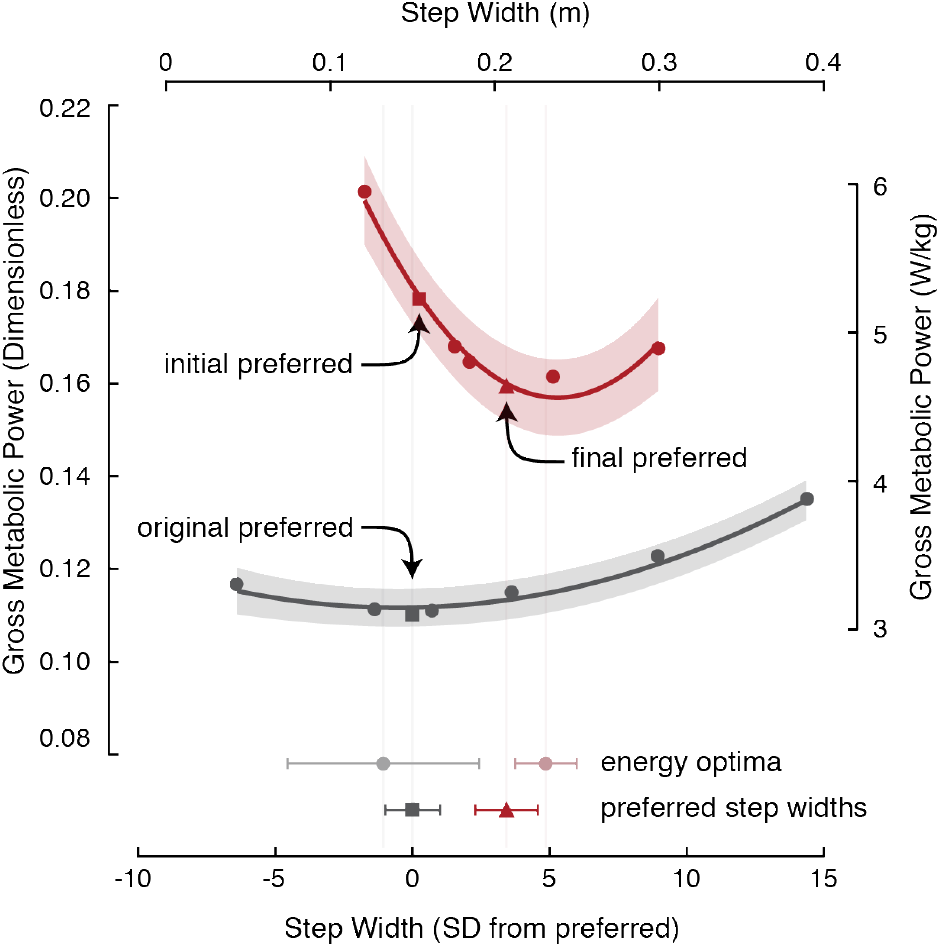
Original cost landscape (grey) from existing data in different subjects(Donelan et al., 2001) and measured new cost landscape (red). In each cost landscape, we averaged measured step width and energetic cost across subjects. We fit second-order polynomial curves to both the original and new cost landscapes. The shading shows their 95% confidence intervals. We calculated the original preferred width (grey square) as the average width across subjects in familiar walking conditions(Donelan et al., 2001), the initial preferred width (red square) as the average width across subjects when the controller is off, and the final preferred width (red triangle) as the average width across subjects when the controller is on and following enforced experience. We calculated the variability in these preferred widths as the standard deviation of the original and final preferred widths across subjects. We calculated the original (grey circle) and new (red circle) energy optimal widths as the average fitted minima across subjects in the original and new cost landscapes. We calculated the variability in these energy optimal widths as the standard deviation of the fitted minima across subjects in the original and new cost landscapes. Error bars represent 1 standard deviation.

We found that subjects needed experience with a lower cost width in the new cost landscape to initiate optimization and adapt towards the new energy optimal width. We measured subjects self-selected widths before giving enforced experience and found that they did not spontaneously adapt towards the new energy optimal width, walking at an average width of −0.7 ± 1.3 SD from initial preferred (*p* = 0.9; Fig. 4b pre). They also returned back to their initial preferred width in self-selected steps after being held at a higher cost width (narrower width) (*p* = 0.2; Fig. 4b n1). Subjects only initiated optimization in self-selected steps after being held at a lower cost width (wider width), where they walked at an average width of 1.6 ± 0.6 SD from initial preferred (p = 9.2×10^−5^; Fig. 4b w1). After optimization was initiated, they gradually converged on the energy optimal width (Fig. 4b, w1-post) with an average time constant of 248 seconds (95% CI [247 249]), or about 468 steps (Fig. 4a). After converging on the final preferred width, which was towards the energy optimal width, subjects quickly returned to it when perturbed away (Fig. 4b, w1-post).

**Fig. 4.**
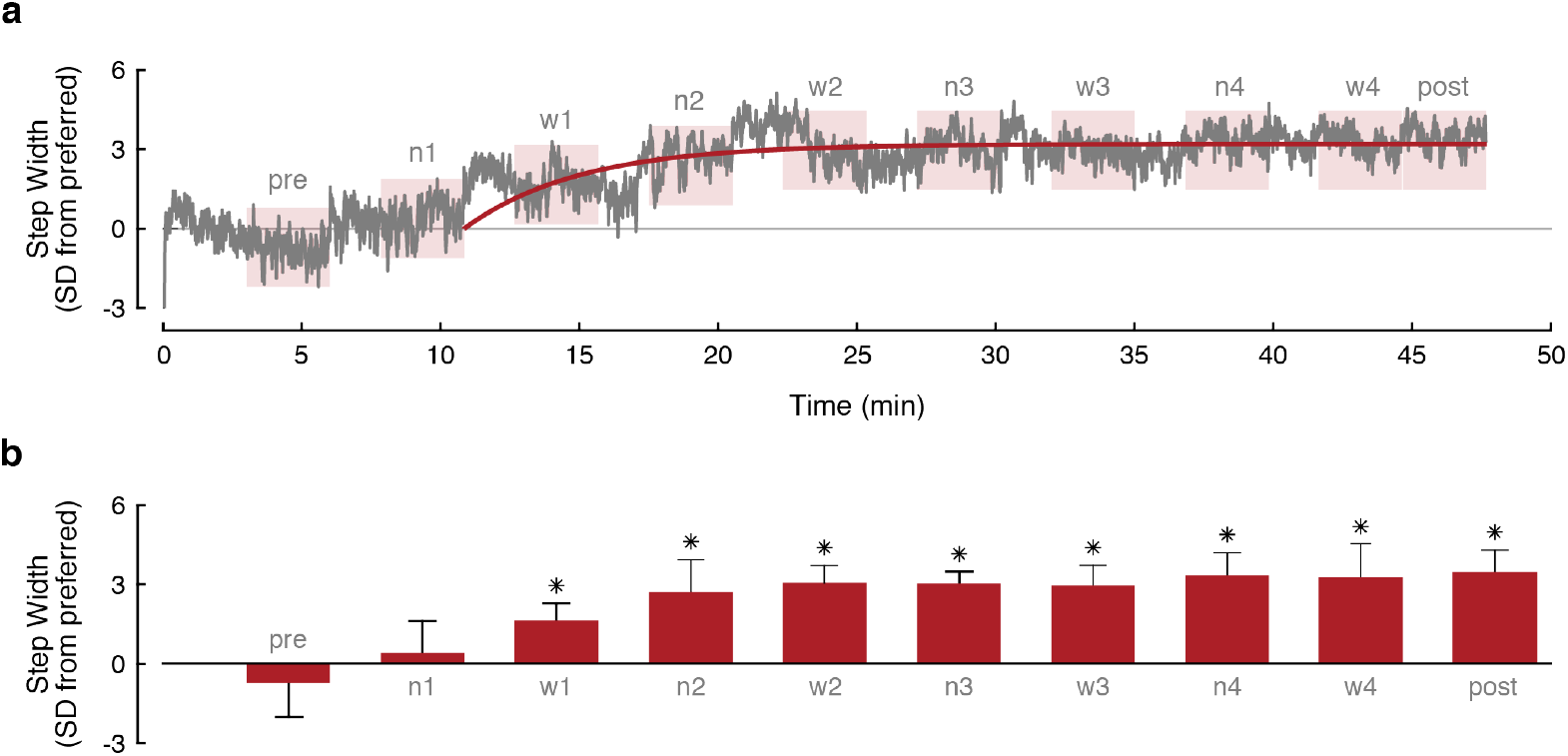
Rate of adaptation. (a) Step width averaged across subjects when allowed to self-select widths prior to enforced experience (pre), during narrow (n1-n4) and wide releases (w1-w4), and following enforced experience (post). The rate of adaptation after optimization was initiated is represented by the time constant of the fitted exponential model. The regions of red shading illustrate the averaging windows. (b) Average widths during period prior to enforced experience, narrow releases, wide releases, and period following enforced experience. Error bars represent 1 standard deviation. Asterisks indicate statistically significant differences in width when compared to initial preferred width.

Subjects adapted their step width to a width that reduced energetic cost. On average, subjects adapted their width 3.5 ± 0.8 SD wider than that initially preferred (*p* = 3.1 x 10^−6^; Fig. 4b post) and reduced energetic cost by 14.4% ± 6.1% relative to the cost of the initial preferred width in the new cost landscape (*p* = 2.4×10^−4^; Fig. 3). This adaptation is not by random chance—a step width 3.3 standard deviations wider than preferred is likely to happen only once in every 1000 steps during normal walking. Although this final preferred width is slightly but significantly narrower than the new energy optimal width (*p* = 2.2 x10^−4^), the costs of these widths are not significantly different (*p* = 0.6).

## DISCUSSION

We tested the hypothesis that energy optimization is a major objective in the real-time control of step width by creating a new cost landscape and then measuring whether people adapt towards the new energy optimal width. Our previous understanding of preferred width was that it is energetically optimal in familiar walking conditions, yet we did not know the timescale by which this preference is established (Donelan et al., 2001). Here we find that the nervous system establishes this preferred width by optimizing energetic cost in real-time.

Our experiment has several limitations. One concern is that people may naturally prefer to walk at a different width when walking at an incline, and that this width is the same as the new energy optimal width. Recent work, as well as our pilot experiments, indicate that width changes minimally within the range of inclines that subjects experience in our study (Kawamura et al., 1991; Sun et al., 1996). Another concern is that subjects are adapting their width to achieve level walking rather than to optimize energy. We partially addressed this by making the minimum incline occur at a non-zero treadmill slope (~4°). A limitation of this design, however, is that the minimum incline is also the energy optimal incline. Thus, we cannot distinguish between a nervous system objective of walking at the shallowest incline from an objective of minimizing energy.

The nervous system s criteria for initiating step width optimization in this experiment, and the process used to converge towards energy optimal widths, is consistent with what we have previously observed for step frequency optimization using knee exoskeletons. The first similarity between studies is that perturbations towards lower cost gaits initiates the optimization process (Selinger et al., 2018; 2015). Experience with this lower cost gait may cue the nervous system to explore the new cost landscape by indicating that its preferred gait is now energetically suboptimal. Without this cue, the nervous system exploits the previously optimal gait, perhaps because an increased cost at the preferred gait does not necessarily imply a change to the cost landscape shape. The second similarity between studies is that after initiating optimization, people converged within minutes towards the energy optimum. Third, the nervous system learned to predict this energetically optimal width, rapidly returning to it when perturbed away (Fig. 4a) (Selinger et al., 2015). These behaviors were common to our two studies, even though we used two different methods of shifting the energy optimal gait and studied two different gait parameters. This suggests that continuous energy optimization is a dominant and general objective of nervous systems in young healthy walking, and that our observed behaviors are common characteristics of the nervous system’s energy optimization.

There are, however, some differences between experiments in how the nervous system initiates optimization as well as the rate at which it converges on new energy optimal gaits. One difference is that no subjects in our current experiment spontaneously optimized their step width, whereas a small subset of people spontaneously optimize step frequency (Selinger et al., 2018; 2015). One possible explanation for this difference is that the general population does have spontaneous step width optimizers, but by chance, we did not have any in our random sample. A more mechanistic candidate explanation is that the nervous system is not primed to identify differences in energetic cost with step width because they are normally relatively small due to the shallowness of the cost landscape around the preferred width. The step width cost landscape varies up to only 0.6% of the cost at preferred step width when walking at widths within ± 3 SD from preferred, whereas the step frequency cost landscape varies up to 1.4% (Donelan et al., 2001; Umberger and Martin, 2007). The second difference is that subjects converged on the energy optimum slower in our step width study compared to our step frequency study. In our step width study, people gradually converged on the energy optimal width with a time constant of 248 seconds (Fig. 4a). Others have found similar rates of adaptation (i.e. hundreds of seconds) in converging on energy optimal movements in both split-belt walking and reaching paradigms (Finley et al., 2013; Huang et al., 2012). However, this is not a general feature of energy optimization—in our step frequency study, people converged on the new energy optimal frequency with a time constant of 11 seconds once optimization was initiated (Selinger et al., 2015). Simple reinforcement learning models predict these different rates of adaptation, ranging from tens to hundreds of seconds, given a change to the learning rate (Selinger et al., 2018). The learning rate determines how past predictions of energetic cost are weighted against new measures of energetic cost. The nervous system may adjust its learning rate in response to a change in confidence in either the predicted energetic cost or the accuracy of the newly measured energetic cost. Given this flexibility, we don ‘ t consider the differences in convergence rates to be evidence against a shared adaptation process.

The nervous system likely determines step width by optimizing energy simultaneously with other objectives. Although the preferred widths are near their energy optimal widths, they do not perfectly coincide. People naturally prefer a width that is slightly wider than the energy optimal width, and in our new cost landscape, they prefer a width that is narrower than the energy optimal width (Donelan et al., 2001). The influence of other objectives may explain these differences. To better understand the importance of other objectives in the nervous system’s control of step width, we modeled an objective function that combined energy with other contributors (Appendix A). There are a number of other objectives that could influence step width, such as stability and maneuverability (Acasio et al., 2017; Dean et al., 2007; Jindrich and Qiao, 2009; McAndrew Young and Dingwell, 2012; Patla et al., 1991; Wu et al., 2015). One way to accommodate not knowing what these non-energy objectives are, or their individual relationships with step width, is to consider their sum as a single additional term in the objective function. With some assumptions, including that the non-energy objective makes the same contribution in both original and new energy cost landscapes, we can estimate its shape using the differences between the preferred widths and the energy optimal widths. Its shape is parabolic, with a minimum at a step width slightly wider than the original preferred (Fig. 5a and b). When compared to the shape of the original energetic cost landscape, its dependence on step width is relatively steep, shifting the minimum of the total objective function, and thus the preferred width, closer to its minimum than that of the energetic cost (Fig. 5a). Their relative contribution to preferred width is not fixed but depends on condition. In our experiment, the new energetic cost landscape is steeper than the original cost landscape, increasing the influence of energetic cost on preferred width in the new cost landscape (Fig. 5b). In everyday walking, the contribution from energy and non-energy objectives to the real-time control of gait will depend not only on biomechanics, but also on how heavily the nervous system weights their individual importance.

**Fig. 5.**
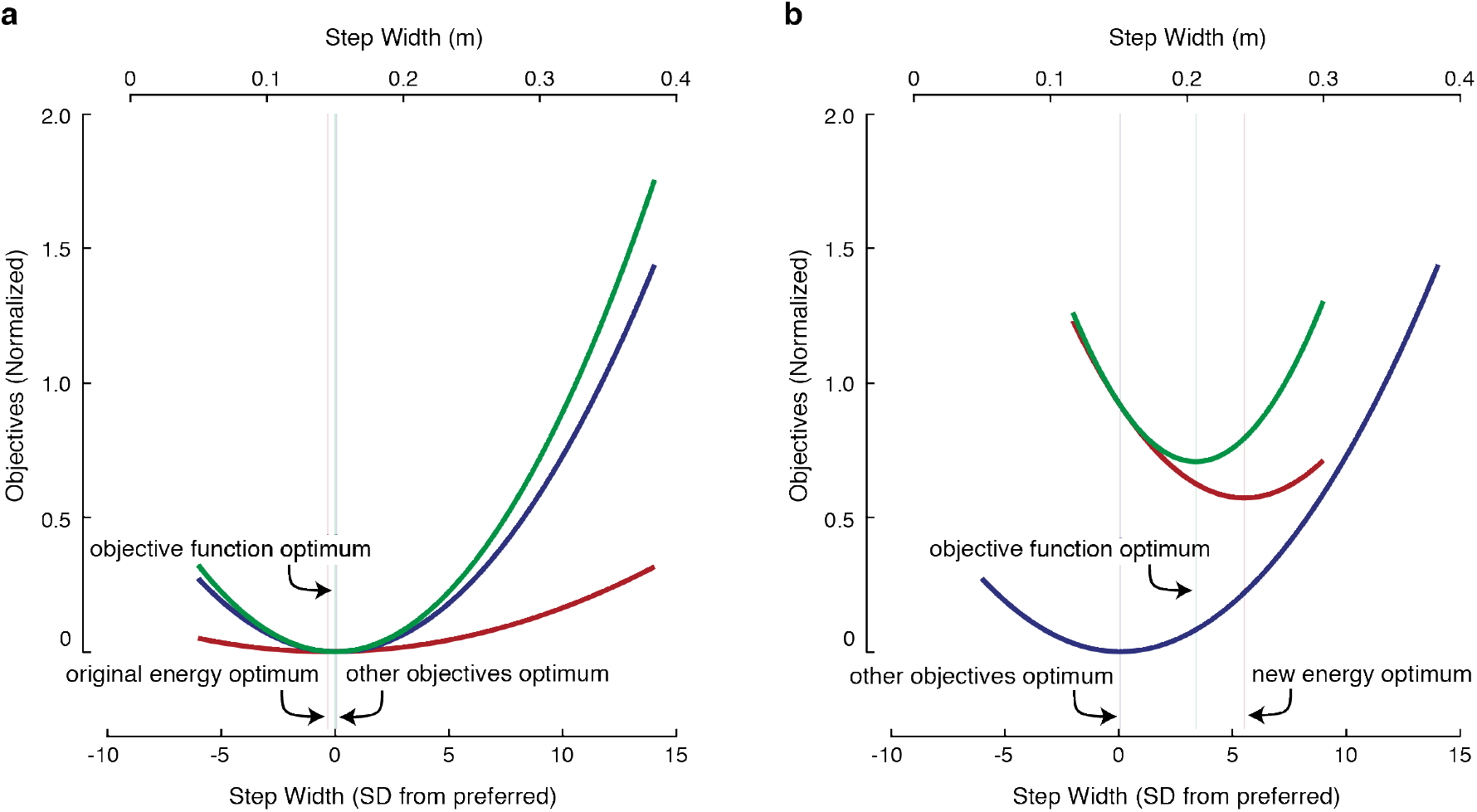
Total objective function for determining preferred step width. We modeled this objective function as the sum of energetic cost and the summed contribution of all other objectives. The total objective function is shown in green, energetic cost shown in red, and contribution of all other objectives shown in blue. To solve for the contribution of all other objectives, we solved for the polynomial coefficients that produced a curve that, when summed with energetic cost, brought the total objective function optimum to (a) the initial preferred width when the controller is off and (b) the final preferred width when the controller is on. Each vertical line represents the location of the optimal step width for either the total objective function, energetic cost, or contribution of other objectives. When the controller is off (a), the vertical lines for the total objective function and contribution of other objectives are nearly coincident.

## ACKNOWLEDGEMENTS

This work was supported a Vanier Canadian Graduate Scholarship to SJA, as well as a Discovery Grant from the Natural Sciences and Engineering Research Council of Canada and a US Army Research Office grant (W911NF-13-1-0268) to JMD. We thank the SFU Locomotion Lab for their constructive feedback on manuscript drafts.

## APPENDIX A

We modeled the nervous system s objective function for determining preferred step width as the sum of energetic cost and the summed contribution of all other objectives:

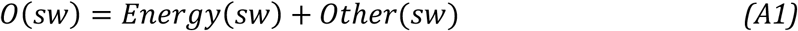

Here, we refer to people walking in the original cost landscape as *controller-off* and people walking in the new cost landscape as *controller-on*. The dependence of energetic cost on step width both within the controller-off and controller-on conditions is well described by a parabola, albeit with different coefficients that depend upon condition:

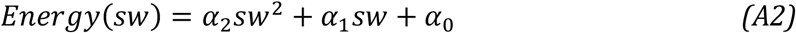

where *α*_2_, *α*_1_, and *α*_0_ are the polynomial coefficients for energetic cost and *sw* is step width. We find the value of these coefficients by fitting a polynomial of this form to our measured energetic cost at different step widths. For presentation purposes, we subtract the minimum cost in the controller-off condition from the energetic cost in both conditions, and then normalize by the controller-off minimum. It is unknown how the other contributors depend on step width. Here we assume that they are also well described by a 2^nd^ order polynomial:

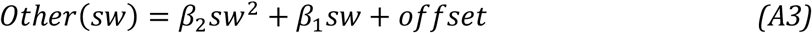

where *β*_2_ and *β*_1_ are unknown coefficients for which we will solve. The purpose of the *offset* term is to force the minimum cost to be zero like the controller-off energy term. It is not a free parameter as its value is uniquely determined by the values of *β*_2_ and *β*_1_. Expanding *Eq. A1* for the two controller conditions yields:

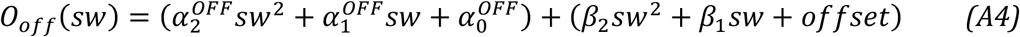

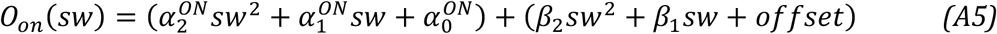

where *O_off_* is the total cost of step width in the controller-off condition and *O_on_* is the total cost of step width in the controller-on condition. Importantly, while the contribution of energetic cost changes with the controller condition, we assume that the summed contribution of the other objectives does not.

We solved for unknowns *β*_2_ and *β*_1_ by first constraining the total cost optimum in the controller-off condition to be at the original preferred step width (0 SD):

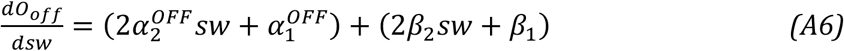

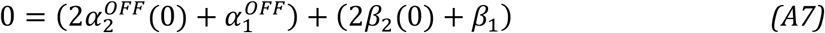

and the total cost optimum in the controller-on condition to be at the final preferred step width (3.4 SD):

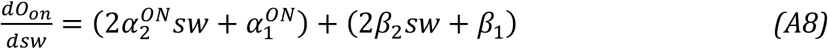

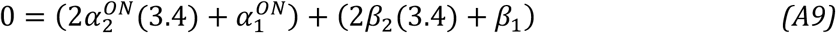

We solved *Eq. A7* and *Eq. A9* simultaneously for unknowns *β*_2_ and *β*_1_. We then substituted in the known values for our coefficients 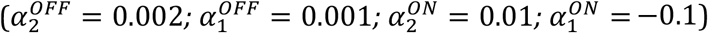 which yielded *β*_2_ = 0.005 and *β*_1_ = −0.001. This parabola has a minimum at 0.1 SD from original preferred.

